# Taxonomic Reassignment of *Pseudohaptolina Birgeri* comb. nov. (Haptophyta)

**DOI:** 10.1101/2020.05.06.081489

**Authors:** Catherine Gérikas Ribeiro, Adriana Lopes dos Santos, Ian Probert, Daniel Vaulot, Bente Edvardsen

## Abstract

The haptophyte genus *Pseudohaptolina* (formerly *Chrysochromulina* clade B1-3) currently harbors two species: *Pseudohaptolina arctica* and *Pseudohap-tolina sorokinii*. In addition, *Chrysochromulina birgeri* is expected to belong to this genus due to its morphological similarity to *P. sorokinii*, but has not yet been genetically characterized. A strain belonging to *Pseudohaptolina* was brought into culture from Arctic waters, characterized by 18S and 28S rRNA gene sequencing as well as optical and transmission electron microscopy, and deposited in the Roscoff Culture Collection with the code RCC5270. Molecular and morphological data from RCC5270 were compared with those from previously described *Pseudohaptolina* and *Pseudohaptolina*-like species. Strain RCC5270 showed strong phylogenetic affinity to *P. sorokinii*, but TEM observations showed that RCC5270 possesses three types of organic body scale, rather than two as originally described in *P. sorokinii*. We found that the occurrence of three scale types is likely to have been overlooked in the original descriptions of both *P. sorokinii* and *C. birgeri*. We also found that environmental metabarcodes identical to the sequence of RCC5270 were abundant in the location from which *C. birgeri* was initially described (Gulf of Finland). We conclude that *P. sorokinii* and *C. birgeri* are conspecific and *P. sorokinii* is therefore synonymous with *C. birgeri*. Based on its phylogenetic placement and nomenclatural priority we propose the new combination *Pseudohaptolina birgeri* and emend the description of this species.

## Introduction

Haptophyte identification is based on both molecular phylogeny and comparison of morphological features such as cell shape, length and movement of the haptonema, and ornamentation of organic body scales. The genus *Pseudohaptolina* was erected from the former *Chrysochromulina* B1-3 clade (Edvardsen et al., 2011). Like most haptophytes, *Pseudohaptolina* are solitary, flagellated and photosynthetic, with two species currently described: the type species *Pseudohaptolina arctica* Edvardsen & Eikrem (Edvardsen et al., 2011) and *Pseudohaptolina sorokinii* Stonik, Efimova & Orlova (Orlova et al., 2016). Both of these *Pseudohaptolina* species were described from high latitude northern hemisphere marine waters, *P. sorokinii* having been collected during an under-ice algal bloom in Amurskiy Bay in the northwestern Sea of Japan (Orlova et al., 2016). A new representative strain from the genus *Pseudohaptolina* was brought into culture from Canadian Arctic waters in 2016 (Gérikas Ribeiro et al., 2020) allowing comparison to previously described *Pseudohaptolina* species using morphological and genetic features.

## Material and Methods

Strain RCC5270 was isolated into clonal culture from Canadian Arctic waters in 2016 (Gérikas Ribeiro et al., 2020), more specifically from Baffin Bay close to the Inuit village of Qikiqtarjuaq, Nunavut on Baffin Island (67^*◦*^28’ N, 63^*◦*^47’ W). The strain was identified by 18S rRNA gene sequencing and optical microscopy and deposited in the Roscoff Culture Collection (http://roscoff-culture-collection.org) with the code RCC5270. Strain RCC5268 was recovered from the same sample than RCC5270 and its 18S rRNA sequence (MH764749) shares 100% similarity of with that of RCC5270.

The nearly complete 18S rRNA gene was amplified using the primers 63F (5’-ACGCTTGTCTCAAAGATTA-3’) and 1818R (5’-ACGGAAACCTTGTTACGA-3’) (Lepère et al., 2011) and sequenced using the same primers and the internal primer 528F (5’-CCGCGGTAATTCCAGCTC-3’) (Zhu et al., 2005). The 28S rRNA gene was amplified and sequenced using primers D1R (5’-ACCCGCTGAATTTAAGCATA-3’) and D3Ca (5’-ACGAACGATTTGCACGTCAG-3’) (Lenaers et al., 1989). Sequencing was performed at Macrogen Europe (https://dna.macrogen-europe.com). Consensus sequences were generated using *de novo* assembly in Geneious® 10 (Kearse et al., 2012). The RCC5270 18S and 28S rRNA gene sequences were deposited in GenBank under accession numbers MT311519 and MT311520, respectively. For phylogenies, sequences from strain RCC5270 were aligned to closely related Haptophyta sequences from Genbank using the Muscle plugin in Geneious® 10 (Kearse et al., 2012).

Samples for transmission electron microscopy (TEM) were prepared as whole mounts fixed with osmium vapor following Eikrem (1996) with slight modifications (cooling of all equipment). Observations were made using a Jeol JEM-2010 FEG at the Imaging Core Facility at the Station Biologique de Roscoff, France. The size of more than 100 scales from RCC5270 and RCC5268 was measured from TEM micrographs using the imaging software ImageJ (https://imagej.nih.gov/ij/). Representative images are available at http://www.roscoff-culture-collection.org/rcc-strain-details/5270.

In order to determine the oceanic distribution of the species corresponding to RCC5270, we examined a large set of publicly available metabarcode datasets (Table 1) covering the V4 and V9 region of the 18S rRNA gene. Twenty-one oceanic 18S rRNA metabarcode datasets were downloaded and reprocessed with the *dada2* R package (Callahan et al., 2016) following the standard operating procedure https://benjjneb.github.io/dada2/tutorial.html in order to produce amplicon single variants (ASVs). The taxonomy of each ASV was assigned using the *dada2* assignTaxonomy function against version 4.12 of the PR^2^ database (Guillou et al., 2013) available at https://github.com/pr2database/pr2database/releases/tag/v4.12.0. Twenty datasets corresponded to the V4 of the 18S rRNA gene, and one to the V9 region (Tara *Oceans*). ASVs with a 100% match to the sequence of RCC5270 were selected and the number of reads in each sample determined using the R library dplyr. Maps and figures were drawn using the R libraries *ggplot2*, *sf* and *cowplot*.

**Table 1.**
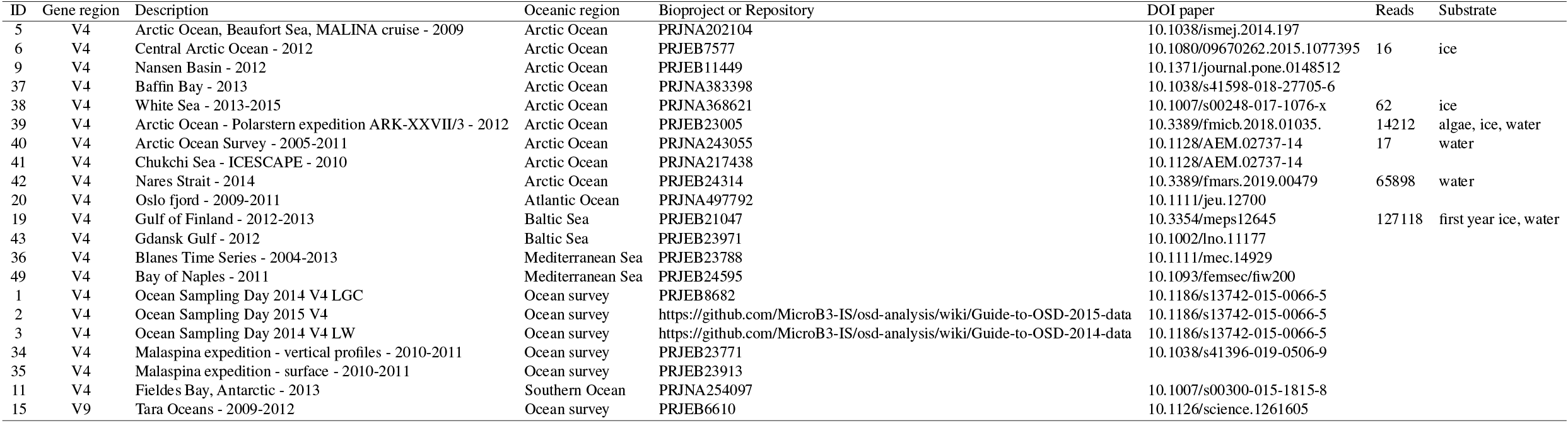
Datasets considered for metabarcode analysis. These 21 datasets correspond to the V4 (20) and V9 (1) regions of the 18S rRNA gene. All datasets have been processed with the dada2 software (Callahan et al., 2016) to extract ASV (amplicon single variants) and assigned using the PR2 database (Guillou et al., 2013).

### Results and Discussion

The 18S rRNA gene sequence from RCC5270 was compared with similar sequences in GenBank including those from previously described *Pseudohaptolina* species. The best match of the sequence was to the two *P. sorokinii* 18S rRNA sequences in GenBank (KF684962 and KU589286), both linked to its original description, although only KF684962 is cited in the text of the original description. The 18S rRNA gene sequence of strain RCC5270 differs from sequence KF684962 by five base pairs (four substitutions and one deletion) in a 1,655 bp alignment and by only one base pair deletion when compared to KU589286 (1,213 bp alignment). The divergences from KF684962 seem to originate from sequencing errors in the *P. sorokinii* description, since they occur in well conserved positions (Figure 1) and when there is a base variation within these positions in related haptophytes, they do not match with those in the *P. sorokinii* sequence (Figure 1). Furthermore, the two sequences linked to the original description of *P. sorokinii* do not share the same substitutions.

**Figure 1.**
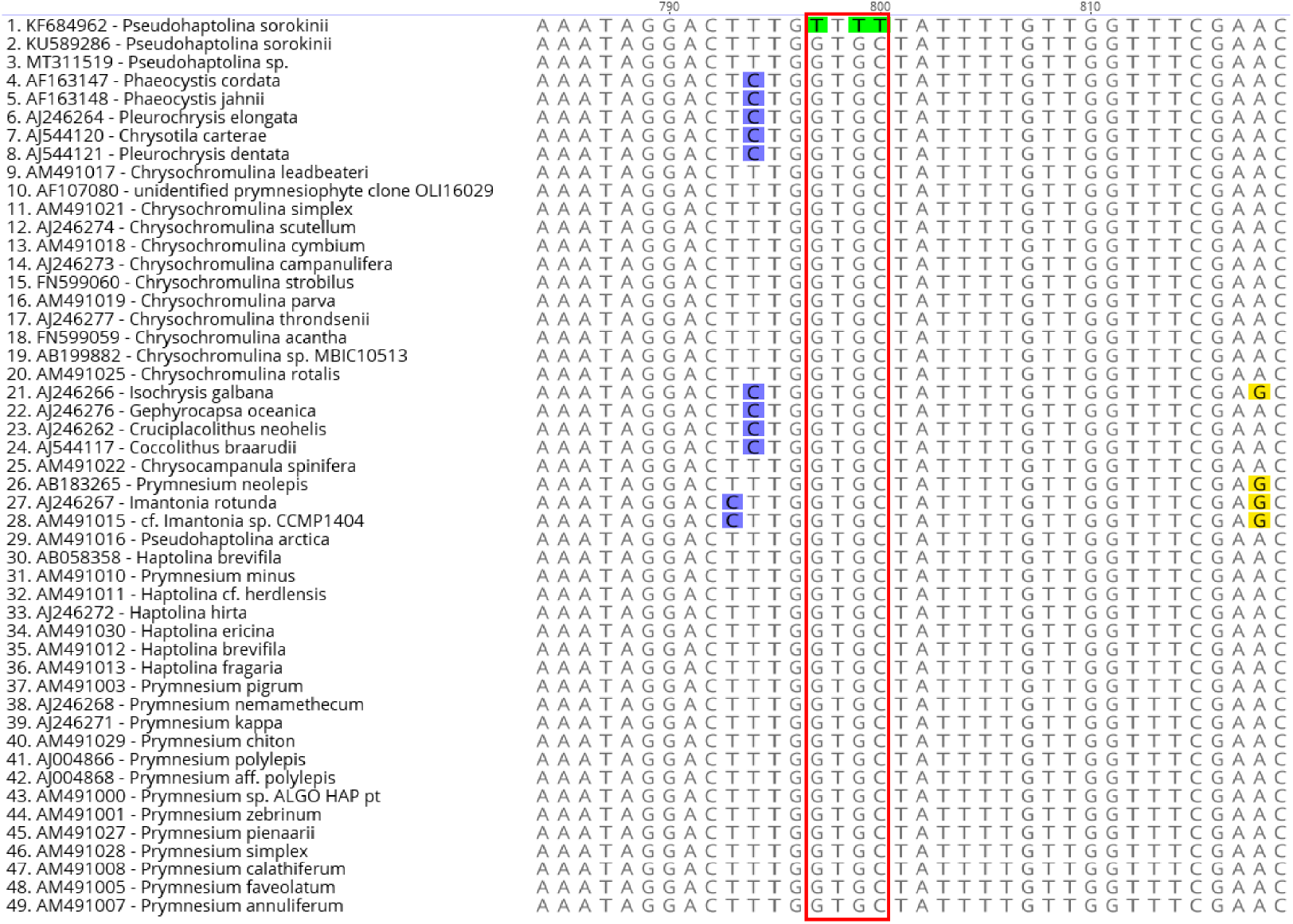
Partial V4 18S rRNA gene sequence alignment showing RCC5270 and *P. sorokinii* KF684962 sequence in comparison to closely related groups; three substitutions are visible in the latter, but they are not shared by any other sequence, including *P. sorokinii* KU589286.

The 28S rRNA gene sequence from RCC5270 has a six base pair difference to the only *P. sorokinii* 28S rRNA sequence available in GenBank (KU589284), which did not originate from the same isolate used for the description of *P. sorokinii*, and is not mentioned in Orlova et al. (2016). Both RCC5270 28S rRNA and KU589284 best hits in GenBank correspond to the environmental clone KU898784 from a sea ice sample in the Barrow Sea (Hassett et al., 2017), with 100% and 98% similarity, respectively.

The shape, size and ornamentation of the organic body scales are taxonomically important characters in Haptophyta, and usually more than one type of body scale occurs per species. *Chrysochromulina birgeri* Hällfors & Niemi (Hällfors and Niemi, 1974) was described before the genus *Pseudohaptolina* was erected, but is expected to be incorporated within *Pseudohaptolina* based on its morphological similarity to members of this genus. The discrimination between *C. birgeri* and other *Pseudohaptolina* species is only possible through morphological examination, since no molecular data or culture strains are available from its first description (Hällfors 1974). *C. birgeri*, *P. arctica* and *P. sorokinii* were all described as possessing two types of body scale (Hällfors and Niemi, 1974; Edvardsen et al., 2011; Orlova et al., 2016), usually referred to as ‘small’ and ‘large’ scales. For the *P. sorokinii* description (Orlova et al., 2016), three morphological features of the organic body scales are indicated as distinctive enough to assign it to a new species: horn morphology, shape of the connecting bridge and density of radial ribs. However, apart from the feature ‘number of radial ribs arranged in quadrants’ present in the so-called small scales, all other measurements overlap to some extent with those recorded for *C. birgeri* (see Table 1 from Orlova et al., 2016, page 511).

In general, the scale morphology of RCC5270 corresponds closely to that described for *C. birgeri* and *P. sorokinii*, including a radial pattern of ribs arranged in quadrants that coincide with the two orthogonal axes of the scale, and two horn-like projections connected by a straight or slightly curved bridge (Figure 2). However, both morphometric data and observations of TEM images of RCC5270 indicate that at least three types of organic scales can be differentiated (Table 2, Figure 2) using scale length, width and distance between the horns, and number of radial ribs per quadrant (Figure 4). Small scales of strain RCC5270 have 37-39 ribs on each quadrant (Figure 2B), as in the description of *C. birgeri* (Hällfors & Niemi, 1974), whereas the medium scales have 54-56 and large scales have 63-68 radial ribs per quadrant (Table 2). The distinction between small and medium scales is, however, most readily visible when comparing scale length *versus* width (Figure 4A). Medium and large scales have somewhat overlapping sizes, so their separation is better achieved by comparing distance between the horn bases *versus* width (Figure 4B), due to a clear distinctive horn bridge structure, with large scales presenting bigger and usually slightly curved bridges (Figure 2).

**Table 2.** Comparison of organic scale measurements between RCC5270, *P. sorokinii* original description (Orlova et al., 2016), *P. sorokinii* independent measurements, and *C. birgeri* original description (Hällfors & Niemi, 1974).

| Measurements | RCC5270 | *P. sorokinii* description | *P. sorokinii* images | *C. birgeri* description |
| --- | --- | --- | --- | --- |
| **Scale length ($\mu\text{m}$)** |  |  |  |  |
| small | 1.1 - 1.4 (1.2 ± 0.1) | 1.6 - 2.0 (1.9 ± 0.03) | 1.67 - 1.73 | 1.5 - 1.7 |
| medium | 1.5 - 2.4 (1.8 ± 0.2) | NA | 1.8 - 2 | NA |
| large | 1.9 - 2.5 (2.2 ± 0.2) | 2.1 - 3.2 (2.6 ± 0.1) | 2.7 - 2.8 | 2.2 - 2.6 |
| **Scale width ($\mu\text{m}$)** |  |  |  |  |
| small | 0.6 - 1 (0.8 ± 0.1) | 1.2 - 1.9 (1.5 ± 0.05) | 0.90 - 0.95 | 1.1 - 1.4 |
| medium | 1.1 - 1.7 (1.3 ± 0.2) | NA | 1.3 - 2 | NA |
| large | 1.1 - 1.8 (1.5 ± 0.2) | 1.6 - 2.3 (1.9 ± 0.1) | 1.5 - 1.61 | 1.7 - 2.1 |
| **Distance between horn bases ($\mu\text{m}$)** |  |  |  |  |
| small | 0.2 - 0.4 (0.3 ± 0.04) | 0.3 - 0.4 (0.4 ± 0.02) | 0.34 - 0.37 | 0.3 - 0.4 |
| medium | 0.3 - 0.6 (0.4 ± 0.1) | NA | 0.4 - 0.5 | NA |
| large | 0.6 - 1.1 (0.7 ± 0.1) | 0.5 - 0.9 (0.7 ± 0.04) | 1.02 - 1.03 | 0.4 - 0.8 |
| **Horn measurements ($\mu\text{m}$)** |  |  |  |  |
| small | 0.1 - 0.2 (0.1 ± 0.1) | 0.2 - 0.4 (0.3 ± 0.02) | 0.26 - 0.3 | 0.1 - 0.2 |
| medium | 0.1 - 0.2 (0.2 ± 0.03) | NA | 0.3 - 0.4 | NA |
| large | 0.2 - 0.6 (0.3 ± 0.1) | 0.5 - 0.9 (0.7 ± 0.03) | 0.7 - 0.8 | 0.2 - 0.6 |
| **Number of ribs per quadrant** |  |  |  |  |
| small | 37 - 39 | 49 - 57 (52.2 ± 0.8) | NA | c. 38 |
| medium | 54 - 56 | NA | 54- 60 | NA |
| large | 63 - 68 | 52 - 64 (57.8 ± 1.5) | NA | 55 - 68 |

**Figure 2.**
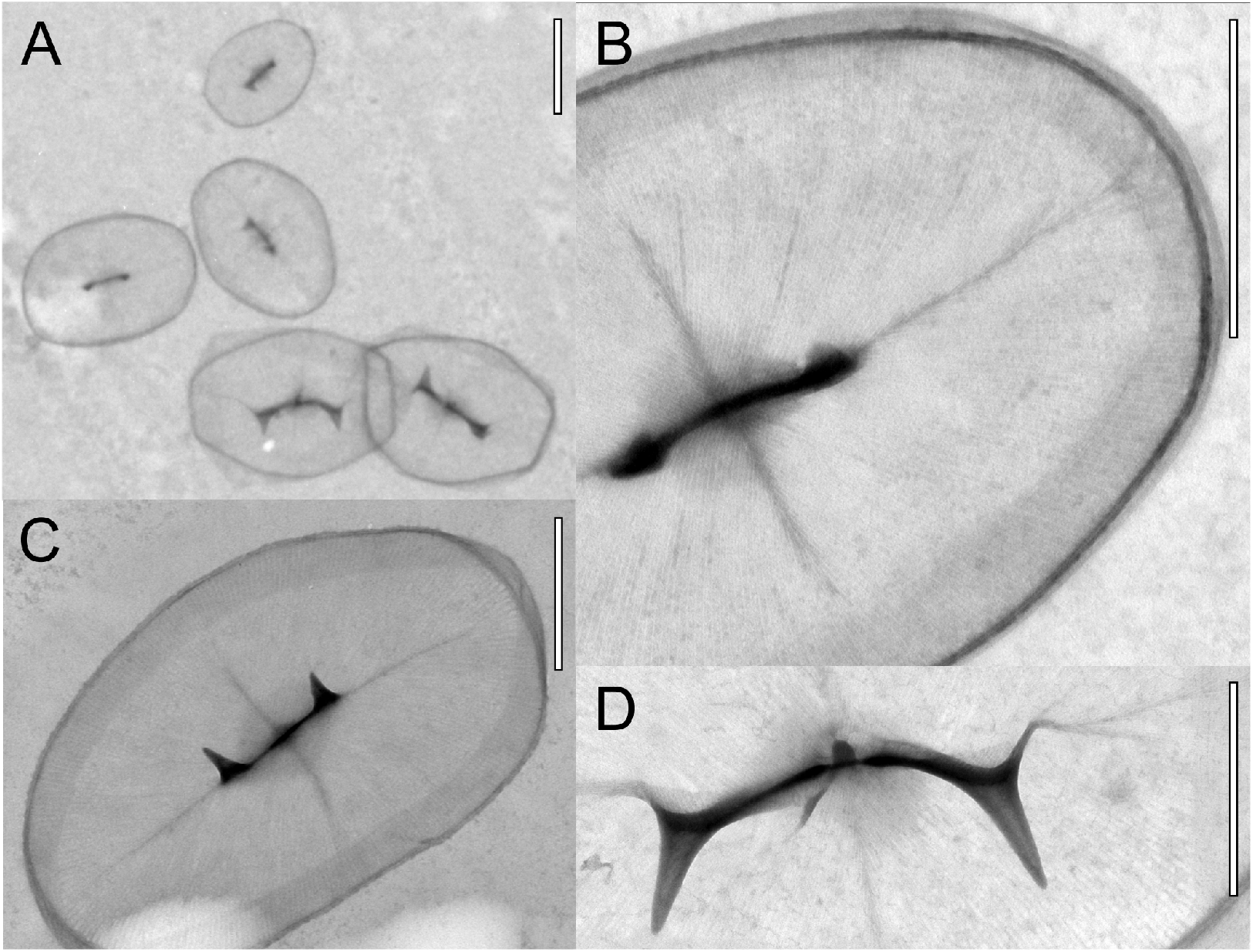
Transmission electron microscopy images of RCC5270. A) Three types of scales: small in the top, medium (short connecting bridges) in the middle and large scales in the bottom. B) Detail of a small scale with approximately 38 ribs in each quadrant. C) medium ellipsoid scale. D) Detail of the slightly curved bridge from a large scale. Scale bars have 1 *μ* m for A and C and 0.5 *μ* m for B and D.

When measurements are conducted on the images displayed in the original descriptions, we found that the three types of scales can be distinguished for *P. sorokinii* (Figure 3A, Figure 4) and most likely also for *C. birgeri*, as shown in Figure 3D. Two *P. sorokinii* organic scales, identified as ‘small scales’ in the original description (Figure 3B and C, see also Orlova et al., 2016, page 510, figures 9 and 11), fall in the same size range as the ‘medium’ scales identified here (Figure 4), which impacts the number of ribs counted. In addition, independent measurements of small scales depicted in figure 8 of the original paper (Figure 3A in the present work), which are true small scales, fall outside the size range of small scales described by Orlova et al. (2016) (Figure 4). Unfortunately, the resolution of available *P. sorokinii* images is not sufficient to perform an independent count of the ribs in the small scales. The size of the connecting bridge was used by Orlova et al. (2016) as a distinctive feature of large scales, so small and medium scales were probably grouped together, which might have led to the discrepancies observed in the number of ribs per quadrant reported in the *P. sorokinii* description. In contrast, in the *C. birgeri* description medium and large scales with evident differences in the connecting bridge structure were grouped together as ‘large’ (Figure 3E and F). It is noteworthy that neither *P. sorokinii* nor RCC5270 scale measurements correspond precisely to the size limits described for *C. birgeri* (Hällfors & Niemi, 1974), particularly for small scales (Figure 4).

**Figure 3.**
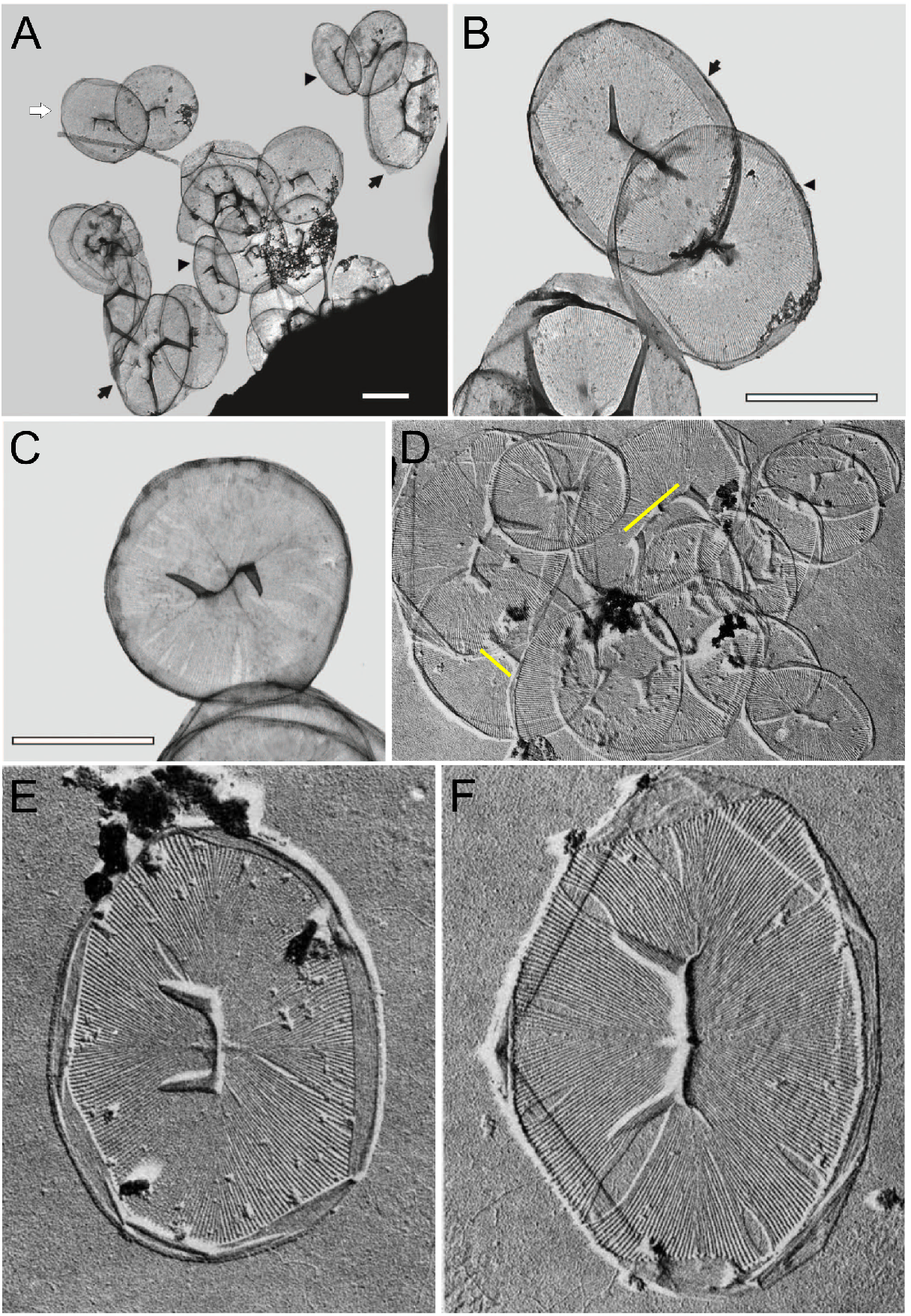
Transmission electron microscopy images for *P. sorokinii* and *C. birgeri* modified from Orlova et al. (2016) and Hällfors & Thomsen (1979), respectively. A) *P. sorokinii* organic body scales showing three types of scales: small (arrow heads), large (arrows) and medium (white arrow, not mentioned in the description paper). B) *P. sorokinii* scales identified as small by Orlova et al. (2016), although its measurements fall within the medium scales size range, being noticeably bigger than the small scales identified in the previous image. C) *P. sorokinii* scale identified as small in the description paper; its round structure, length and width are similar to medium scales. D) *C. birgeri* image with yellow lines highlighting differences in the connecting bridge between the horn bases, the main feature used to distinguish large from medium scales in the present study. For comparison regarding size, one small scale can be seen in the upper right corner of the image. E) *C. birgeri* identified as large by Hällfors & Thomsen (1979) although its features, including the number of ribs per quadrant (54), would correspond to a medium size scale. F) *C. birgeri* true large scale, with a longer distance between horn bases and a slight curved bridge. Scale bars correspond to 1 *μ* m for *P. sorokinii* and no scale bar is available for *C. birgeri*.

**Figure 4.**
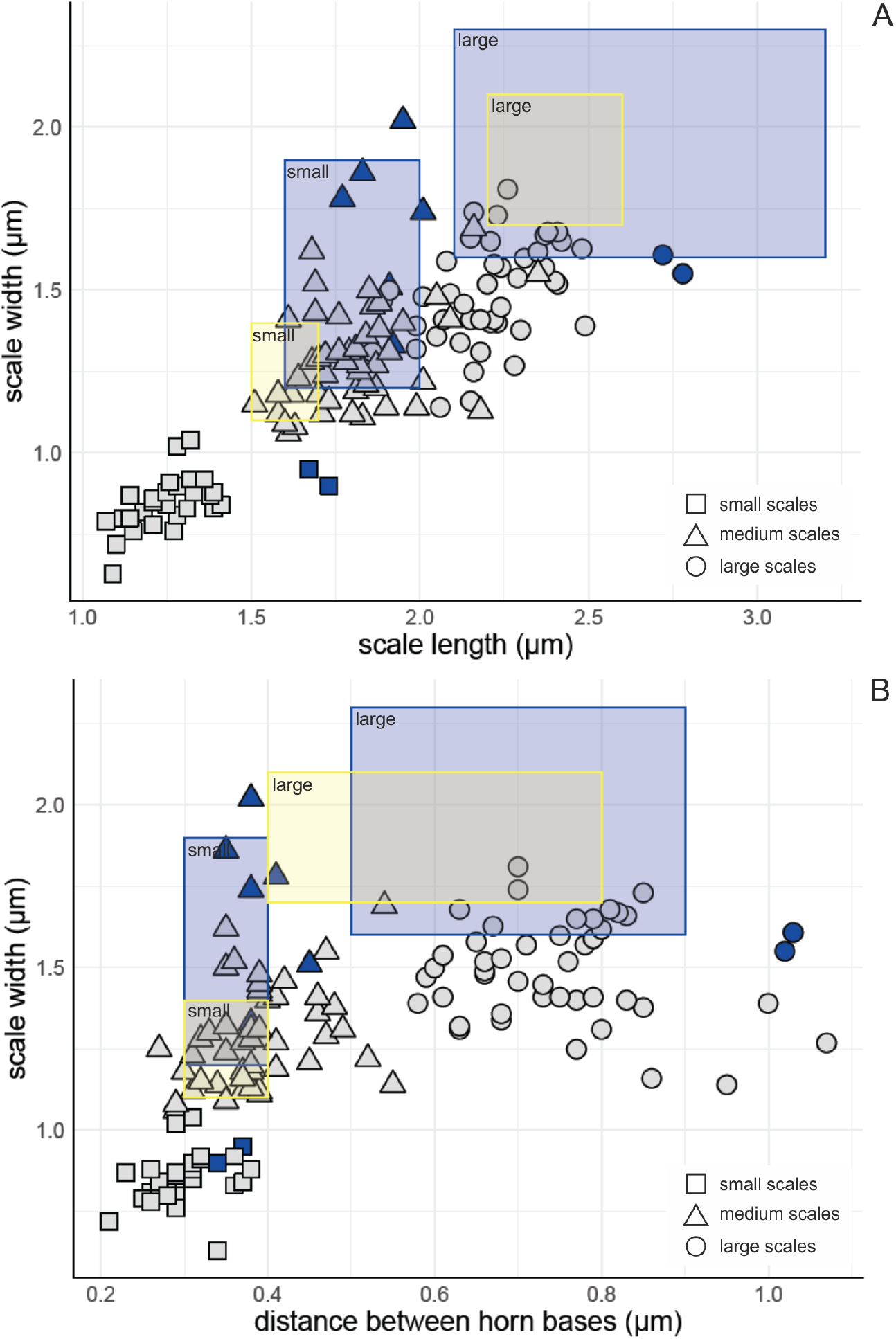
Organic scales measurements from RCC5270 (in grey) and *P. sorokinii* independent measurements from images displayed in Orlova et al. (2016) (in blue). A) Scale length versus width; B) Scale length versus the distance between the horns. Scales visually identified as small scales are represented by squares, medium scales by triangles and large scales by circles. The size limits for *C. birgeri* described in Hällfors & Niemi (1974) and for *P. sorokinii* in Orlova et al. (2016) are displayed as yellow and blue boxes, respectively.

Other morphological characteristics used to differentiate *P. sorokinii* from *C. birgeri* by Orlova et al. (2016) are horn length and the shape of the connecting bridge. Orlova et al. (2016) reported long horn projections and curved connecting bridges, in contrast to the description of *C. birgeri*, although long horn-like projections connected by a curved bridge in large scales have already been reported for *C. birgeri* (Takahashi, 1981; Hällfors and Thomsen, 1979). The horn projections of large scales of RCC5270 are in general smaller than observed by Orlova et al. (2016), but are somewhat superimposed within their size range (Table 2). We also observed curved connecting bridges in the large scales (Figure 2A). There is therefore considerable overlap but some variability in the size and features of scales of RCC5270, *P. sorokinii* and *C. birgeri* which might reflect morphological plasticity within a single species, since heteromorphic life cycles have been observed within the Prymnesiales (Paasche et al., 1990; Edvardsen and Vaulot, 1996).

The metabarcode datasets used to determine the oceanic distribution of RCC5270 correspond to more than 2,200 samples included in large scale surveys such as Ocean Sampling Day (OSD) and the Tara *Oceans* and Malaspina expeditions that sampled a wide range of coastal and oceanic waters as well as more limited studies from polar waters and the Baltic Sea. We did not retrieve any V9 metabarcodes identical to the RCC5270 sequence. We did, however, retrieve six V4 metabarcodes (ASVs) that were 100% identical to the RCC5270 sequence (Figure S1). In contrast, no exact match was found to either KF684962 or KU589286 *P. sorokinii* in any of these datasets, which further corroborates the assumption that the mismatch between 18S rRNA *P. sorokinii* and RCC5270 sequences are due to sequencing errors. The RCC5270 metabarcodes were only observed in the Arctic Ocean and in the Baltic Sea from ice and water samples as well from algal aggregates collected from the deep-sea floor (Figure 5A-B). Metabarcodes identical to the sequence of RCC5270 were particularly abundant in three datasets (Table 1) from the Polarstern expedition in the Central Arctic Ocean (Rapp et al., 2018), from the Nares strait, the northernmost outflow gateway of Baffin Bay (Kalenitchenko et al., 2019) and from the Gulf of Finland (Baltic Sea) (Enberg et al., 2018). At the latter location, which corresponds to the region from which *C. birgeri* was initially described, metabarcodes identical to the RCC5270 sequence first appeared in February in the ice where they peaked in early March and then increased massively in the water column one month later, representing up to 70% of the metabarcodes at the time the ice melted in mid-April (Figure 5C). These data indicate that RCC5270 is an ice alga that can seed and proliferate in the water column and even accumulate on the deep-sea floor.

**Figure 5.**
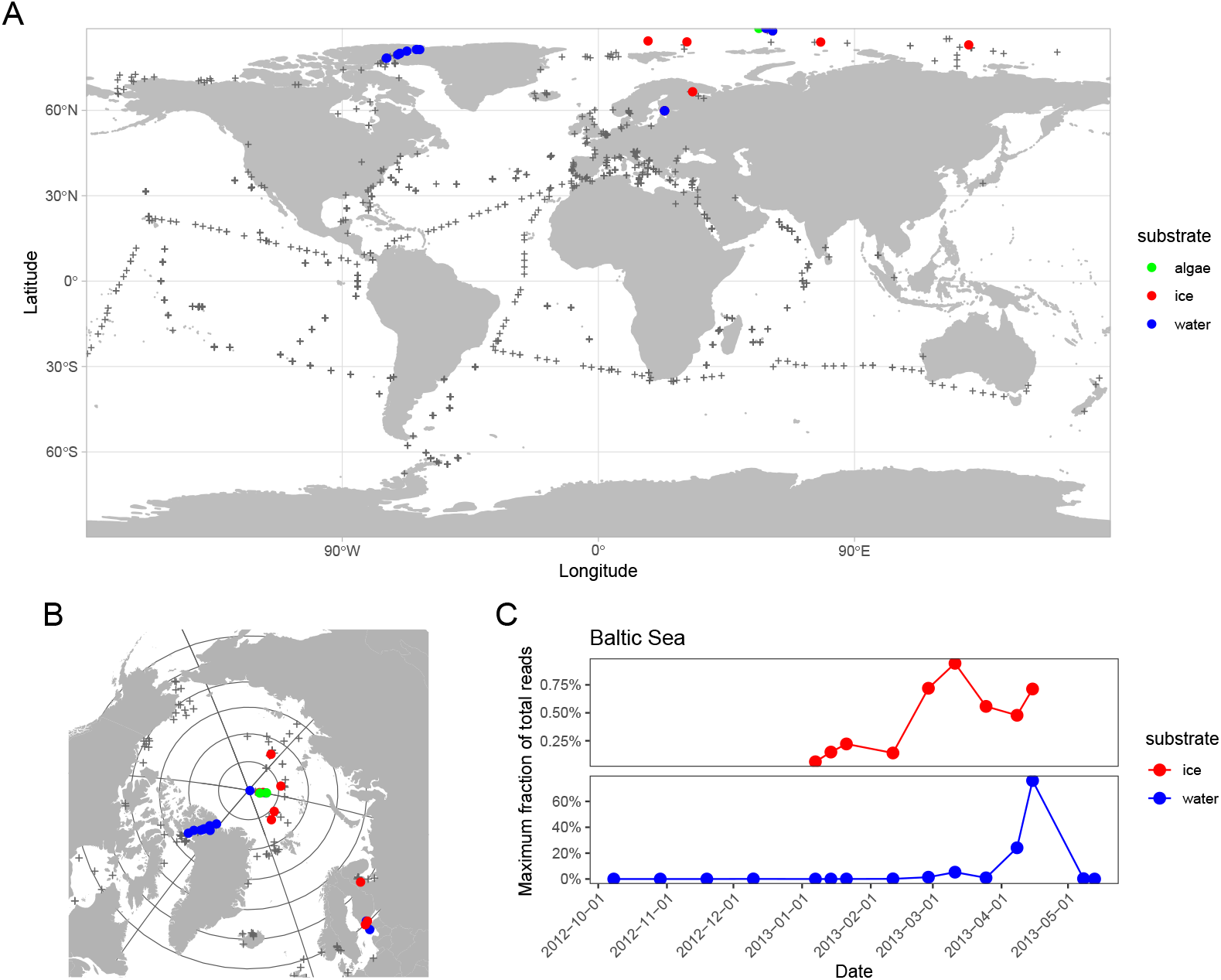
RCC5270 metabarcodes. A) Localisation of stations where 18S rRNA metabarcodes 100% identical to RCC5270 sequence have been detected in public sequence datasets (see Table 1). Color corresponds to substrate. The location of samples where these metabarcodes have not been detected are marked by grey crosses. B) Zoom on the North Pole region. C) Maximum fraction of RCC5270 metabarcodes (excluding Metazoa) as a function of date in the Gulf of Finland (Baltic Sea) in ice and water (Enberg et al., 2018).

## Conclusions

We isolated a culture strain from the Arctic which was genetically affiliated to *P. sorokinii*. Morphological data indicate that a third scale type was overlooked in the original description of *P. sorokinii* (Orlova et al., 2016), impacting the number of radiating ribs described for each scale type. We also found that *C. birgeri* cells have three types of organic body scale, not two as reported in the original description (Hällfors and Niemi, 1974). Metabarcode data indicates that sequences identical to that of RCC5270 were abundant near the type locality of *C. birgerii*. We conclude that *P. sorokinii* is conspecific with the formerly described *C. birgeri* and we therefore transfer *C. birgeri* to the genus *Pseudohaptolina* and emend its description. *P. birgeri* is the valid name for this species due to nomenclatural priority over *P. sorokinii*.

## Taxonomic appendix

*Pseudohaptolina birgeri* (Hällfors & Niemi) Ribeiro and Edvardsen comb. nov. emend. Ribeiro and Edvardsen

### BASIONYM

*Chrysochromulina birgeri* Hällfors & Niemi in Hällfors & Niemi (1974). Memoranda Societatis pro Fauna et Flora Fennica 50. Drawing Fig. 4.

### SYNONYM

*Pseudohaptolina sorokinii* Stonik, Efimova & Orlova.

### EMENDED DESCRIPTION

Scaly covering composed of three round to oval scale types. Small scales have width x length c. 0.6-1.4 x 1.1-1.7, medium scales c. 1.1-2 x 1.5-2.4 and large scales c. 1.1-2.1 x 1.9-2.8 nm. All scales with radial ribs on both distal and proximal faces. Small scales have 37-39 radial ribs per quadrant, medium scales 54-60 and large scales 63-68. Medium and large scales have two horns on the distal face. The distance and form of the horns are different in medium and large scales.

## Contributions

Contributed to conception and design: CGR, IP, DV, BE

Contributed to acquisition of data: CGR, ALS, IP, DV, BE

Contributed to analysis and interpretation of data: CGR, ALS, IP, DV, BE

Drafted and/or revised the article: CGR, ALS, IP, DV, BE

Approved the submitted version for publication: CGR, ALS, IP, DV, BE

## Acknowledgments

We are grateful to Sophie Le Panse from the Merimage microscopy platform at the Roscoff Marine Station for assistance with the transmission electron micrographs and to the Roscoff Culture Collection for maintenance of the algal strain.

## Data accessibility statement

Supporting data have been deposited to GitHub https://github.com/vaulot/Paper-2020-Ribeiro-Pseudohaptolina.

## Supplementary material

### Supplementary Data

Supplementary data are available on GitHub at https://github.com/vaulot/Paper-2020-Ribeiro-Pseudohaptolina

- **Supplementary Data S1**: Alignment of sequences for 18S rRNA gene (fasta file).
- **Supplementary Data S2**: Alignment of sequences for 28S (fasta file).
- **Supplementary Data S3**: Scale measurements (xlsx file).
- **Supplementary Data S4**: Number of *P. sorokinii* reads in each of the metabarcode samples analyzed (xlsx file).
- **Supplementary Data S5**: Alignment of the V4 region of the 18S rRNA for *Pseudohaptolina* reference sequences and metabarcodes (fasta file).

## Supplementary Figures

**Figure S1.**
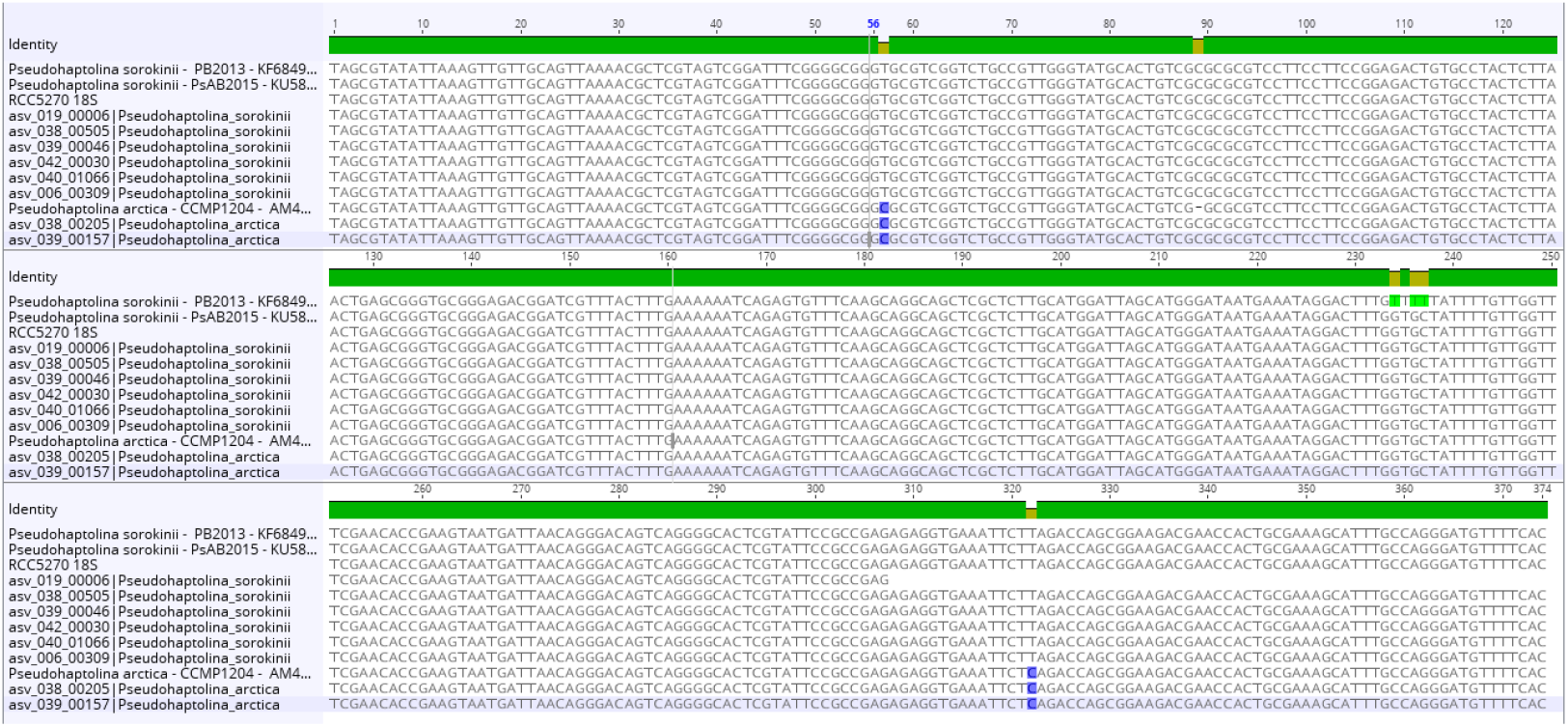
Partial 18S rRNA gene sequence alignment showing RCC5270 and *P. sorokinii* sequences with the *P. sorokinii* metabarcodes identified in the public datasets analyzed (Table 1).

## References

Callahan BJ, McMurdie PJ, Rosen MJ, Han AW, Johnson AJA, et al. 2016. DADA2: High-resolution sample inference from Illumina amplicon data. Nature Methods 13(7): 581–583. doi:10.1038/nmeth.3869.

Edvardsen B, Eikrem W, Throndsen J, Sáez AG, Probert I, et al. 2011. Ribosomal DNA phylogenies and a morphological revision provide the basis for a revised taxonomy of the Prymnesiales (Haptophyta). European Journal of Phycology 46(3): 202–228. doi:10.1080/09670262.2011.594095.

Edvardsen B, Vaulot D. 1996. Ploidy analysis of the two motile forms of *Chrysochromulina polylepis* (Prymnesiophyceae). Journal of Phycology 32: 94–102.

Eikrem W. 1996. *Chrysochromulina throndsenii* sp. nov. (Prymnesiophyceae). Description of a new haptophyte flagellate from Norwegian waters. Phycologia 35(5): 377–380.

Enberg S, Majaneva M, Autio R, Blomster J, Rintala JM. 2018. Phases of mi-croalgal succession in sea ice and the water column in the Baltic Sea from autumn to spring. Marine Ecology Progress Series 599: 19–34. doi: 10.3354/meps12645.

Gérikas Ribeiro C, Lopes dos Santos A, Marie D, Le Gall F, Probert I, et al. 2020. Culturable diversity of Arctic phytoplankton during pack ice melting. Elementa: Science of the Anthropocene 8: 6. doi:10.1101/642264v1.

Guillou L, Bachar D, Audic S, Bass D, Berney C, et al. 2013. The Protist Ribosomal Reference database (PR^2^): a catalog of unicellular eukaryote Small Sub-Unit rRNA sequences with curated taxonomy. Nucleic Acids Research 41(D1): D597–D604. doi:10.1093/nar/gks1160.

Hällfors G, Niemi A. 1974. *Chrysochromulina* (Haptophyceae) bloom under the ice in the Tvarminne Archipelago, southern coast of Finland. Memoranda Societatis pro Fauna et Flora Fennica 50: 89–104.

Hällfors G, Thomsen HA. 1979. Further observations on Chrysochromulina birgeri (Prymnesiophyceae) from the Tvärminne archipelago, SW coast of Finland. Acta Bot Fennica 110(July): 41–46.

Hassett BT, Ducluzeau ALL, Collins RE, Gradinger R. 2017. Spatial distribution of aquatic marine fungi across the western Arctic and sub-Arctic. Environmental Microbiology 19(2): 475–484. doi:10.1111/1462-2920.13371.

Kalenitchenko D, Joli N, Potvin M, Tremblay JÉ, Lovejoy C. 2019. Biodiversity and species change in the Arctic Ocean: A view through the lens of Nares Strait. Frontiers in Marine Science 6(August): 1–17. doi:10.3389/fmars.2019.00479.

Kearse M, Moir R, Wilson A, Stones-Havas S, Cheung M, et al. 2012. Geneious Basic: An integrated and extendable desktop software platform for the organization and analysis of sequence data. Bioinformatics 28(12): 1647–1649. doi:10.1093/bioinformatics/bts199.

Lenaers G, Maroteaux L, Michot B, Herzog M. 1989. Dinoflagellates in evolution. A molecular phylogenetic analysis of large subunit ribosomal RNA. Journal of Molecular Evolution 29(1): 40–51.

Lepère C, Demura M, Kawachi M, Romac S, Probert I, et al. 2011. Wholegenome amplification (WGA) of marine photosynthetic eukaryote populations. FEMS Microbiology Ecology 76: 513–523. doi:10.1111/j.1574-6941.2011.01072.x.

Orlova TY, Efimova KV, Stonik IV. 2016. Morphology and molecular phylogeny of *Pseudohaptolina sorokinii* sp . nov . (Prymnesiales, Haptophyta) from the Sea of Japan, Russia. Phycologia 55(5): 506–514. doi:10.2216/15-107.1.

Paasche E, Edvardsen B, Eikrem W. 1990. A possible alternate stage in the life cycle of *Chrysochromulina polylepis* Manton et Parke (Prymnesiophyceae). Nova Hedwigia Beiheft 100(May 2014): 91–99.

Rapp JZ, Fernández-Méndez M, Bienhold C, Boetius A. 2018. Effects of ice-algal aggregate export on the connectivity of bacterial communities in the central Arctic Ocean. Frontiers in Microbiology 9(May): 1035. doi:10.3389/fmicb.2018.01035.

Takahashi E. 1981. Floristic study of ice algae in the sea ice of a lagoon, Lake Saroma, Hokkaido, Japan. Memoirs of the National Institute of Polar Research 34: 49–56.

Zhu F, Massana R, Not F, Marie D, Vaulot D. 2005. Mapping of picoeucaryotes in marine ecosystems with quantitative PCR of the 18S rRNA gene. FEMS Microbiology Ecology 52: 79–92. doi:10.1016/j.femsec.2004.10.006.

